# The role of FraI in cell-cell communication and differentiation in the hormogonia-forming cyanobacterium Nostoc punctiforme

**DOI:** 10.1101/2024.04.10.588879

**Authors:** Ana Janović, Iris Maldener, Claudia Menzel, Gabriel A. Parrett, Douglas D. Risser

**Affiliations:** Interfaculty Institute of Microbiology and Infection Medicine Tübingen, Organismic Interactions, University of Tübingen, Auf der Morgenstelle 28, 72076 Tübingen, Germany; Department of Biology, University of Colorado, Colorado Springs, USA

**Keywords:** Cell-cell communication, hormogonia, filamentous cyanobacteria, *Nostoc punctiforme*, motility, septal junctions, NpF4142, FraI

## Abstract

Multicellular cyanobacteria, like *Nostoc punctiforme*, rely on septal junctions for cell-cell communication, which is crucial for coordinating various physiological processes including differentiation of N_2_-fixing heterocysts, spore-like akinetes and hormogonia - short, motile filaments important for dispersal. In this study we functionally characterize a protein, encoded by gene *NpF4142*, which in a random mutagenesis approach, initially showed a motility-related function. The reconstructed *NpF4142* knockout mutant exhibits further distinct phenotypic traits, including altered hormogonia formation with significant reduced motility, inability to differentiate heterocysts and filament fragmentation. For that reason, we named the protein FraI (fragmentation phenotype). The mutant displays severely impaired cell-cell communication, due to almost complete absence of the nanopore array in the septal cell wall, which is an essential part of the septal junctions. Despite lack of communication, hormogonia in the Δ*fraI* mutant maintain motility and phototactic behaviour, even though less pronounced than the wild type. This suggests an alternative mechanism for coordinated movement beyond septal junctions. Our study underscores the significance of FraI in nanopore formation and cell differentiation and provides additional evidence for the importance of septal junction formation and communication in various differentiation traits of cyanobacteria. The findings contribute to a deeper understanding of the regulatory networks governing multicellular cyanobacterial behaviour, with implications for broader insights into microbial multicellularity.

**Importance:** The filament-forming cyanobacterium *Nostoc punctiforme* serves as a valuable model for studying cell differentiation, including the formation of nitrogen-fixing heterocysts and hormogonia. Hormogonia filaments play a crucial role in dispersal and plant colonization, providing a nitrogen source through atmospheric nitrogen fixation, thus holding promise for fertilizer-free agriculture. The coordination among the hormogonia cells enabling uniform movement towards the positive signal remains poorly understood. This study investigates the role of septal junction-mediated communication in hormogonia differentiation and motility, by studying a Δ*fraI* mutant with significantly impaired communication. Surprisingly, impaired communication does not abolish synchronized filament movement, suggesting an alternative coordination mechanism. These findings deepen our understanding of cyanobacterial biology and have broader implications for multicellular behavior in prokaryotes.

## Introduction

The filamentous cyanobacterium *Nostoc punctiforme* ATCC 29133/PCC73102 is a true multicellular bacterium, able to form specialized cells which enable the bacterium to adapt to various environmental conditions. Heterocysts are nitrogen-fixing cells that differentiate from semi-regularly positioned vegetative cells upon nitrogen starvation to provide a microoxic environment for the oxygen-labile nitrogenase^1,2^. Hormogonia (see below) are short filaments specialized as motile units allowing dispersal. Finally, unicellular akinetes form under various stress conditions enabling survival as spore-like resistant cells ^3–5^.

Within a filament, cells interact through a remarkable cell-cell communication system, involving so called septal junctions. These multi-protein complexes connect the individual cells by traversing the septal peptidoglycan layer through nanopores^6,7^. On each cytoplasmic side, every septal junction contains a flexible cap and plug structure by which the intercellular connection closes in a reversible manner. This gating mechanism of halting communication is crucial under stress conditions, such as UV-C light exposure, cytoplasm leakage due to filament breakage, or the isolation of senescent cells from the rest of the filament ^7–9^. Mutants with impaired cell-cell communication also exhibit deficiencies in heterocyst development^1,10– 14^, emphasizing the significance of intercellular signaling in cellular differentiation ^1,2,10–15^.

Formation of septal nanopores requires the activity of cell wall amidases. In unicellular bacteria amidases play a pivotal role in cell division, by cleaving the newly synthesized peptidoglycan to facilitate separation of daughter cells ^16^. In multicellular cyanobacteria, cell wall amidases play a role in cell-cell communication, by perforating the septal peptidoglycan^13^. Several amidases were identified in multicellular cyanobacteria ^12,13,17,18^. A knockout mutant of the AmiC2 enzyme in *N. punctiforme* showed severe impairment in cell division, filament morphology and cell-cell communication, caused by the complete absence of nanopores in the septal peptidoglycan ^17^. The *amiC2* mutant is unable to differentiate heterocysts, hormogonia and akinetes^12,17^. The AmiC2 protein primarily localizes along the septal plane of nascent septa and is typically absent in fully developed septa ^12^. AmiC1, another amidase encoded in the same gene operon, is likely essential for *Nostoc punctiforme*, since the *knockout* of this gene was reported to be impossible to create ^12,17^. In the closely related cyanobacterium *Nostoc* sp. PCC 7120 mutants in both orthologues, *amiC1* and *amiC2* as well as double *amiC1amiC2* knockouts were created. All mutants exhibited aberrant nanopore shape and numbers and lower rates of cell-cell communication compared to the wild type ^13^. Furthermore, overexpression of AmiC1 in *Nostoc* sp. 7120 resulted in a lethal phenotype, in contrast to the AmiC2 overexpression, which has no significant effect on growth. The LytM factor Alr3353 is an activator of AmiC1 of *Nostoc* sp. 7120 and its absence leads to a reduction in nanopore numbers and the rate of cell-cell communication ^19^. To date, no other regulators of amidase’s nanopore-drilling activity have been identified.

Transiently formed hormogonia, 10-20 cell-long filaments of *Nostoc punctiforme*, play a crucial role in dispersal and symbiotic plant colonization ^20^. After colonization of the host plant, the hormogonia develop back to vegetative filaments containing heterocysts for N_2_ fixation. This symbiosis is of potential agricultural importance as it enables fertilizer-free cultivation providing nitrogen sources for the crop, as shown for *Oryza sativa* ^21^. Induction of hormogonia differentiation from vegetative filaments can be triggered by various signals such as nitrogen stepdown, red-light exposure or HIF (*hormogonium-inducing factor*) exposure. Following the induction signal, vegetative cells differentiate into hormogonia within 18-24 hours through rapid cell divisions not accompanied by overall biomass growth. These filaments demonstrate photo and chemo-tactic movement, facilitated by a type IV pili machinery on their outer cell’s surface. The movement is additionally supported by the export of an exopolysaccharide sheath known as hormogonia polysaccharide (HPS) ^22^. Directionality is determined by the presence of the HmpF protein at one pole of each cell, positioned uniformly at the leading poles of each cell in motile filaments. Repositioning of the HmpF protein occurs rapidly and synchronously along the filament, typically within about 30 seconds, in response to changes in light conditions, leading to a change in the filament’s course. ^23,24^. A putative amidase encoded by *the tftA* gene was shown to be important for the tapered cell phenotype at the filament’s termini, likely by the degradation of the terminal peptidoglycan. Further, a *tftA* knockout showed reduced migration ability through agar ^25^. However, the role of cell-cell communication in the synchronized movement of the entire filament and during the change of movement’s direction in hormogonia remained uninvestigated. In this study, we identified a mutant lacking the NpF4142 protein, herein designated FraI, that presents decreased hormogonia motility. Characterization of this mutant strain allowed us to show that FraI plays a key role in nanopore formation and hormogonia differentiation. Furthermore, this study showed that transmission of the signal for direction of movement in hormogonia probably does not involve communication through the septal junctions.

## Material and methods

### Cultivation of bacterial strains

As standard medium, *N. punctiforme* and the mutant strains were cultivated in AA/4 medium according to Allen and Arnon ^26^ in a fourfold dilution. To supply a source of fixed nitrogen, the medium was supplemented with 2.5 mM NH_4_Cl and 5 mM MOPS buffer, pH 7.8. In experiments performed in Tübingen the cyanobacteria were cultivated in either liquid BG11 medium ^27^ or AA/4 medium supplemented with 2.5 mM KNO_3_ and 2.5 mM NaNO_3._ Liquid cultures were shaken at 120 rpm, agar plates contained 1% Difco agar. Growth occurred at 28 °C with constant illumination at 25-40 μE m^-2^ s^-1^ Sucralose was added to suppress hormogonia development in concentrations mentioned in the text. Neomycin was used at a concentration of 50 μg/ml in appropriate strains.

### Plasmid and strain construction

For a detailed description of the plasmids, strains, and oligonucleotides used in this study refer to supplemental material, Tables S1. All constructs were sequenced to insure fidelity.

To construct plasmid pDDR558 for in-frame deletion of *fraI* (Npun_F4142), approximately 900 bp of flanking DNA on either side of the gene and several codons at the beginning and end of the gene were amplified via overlap extension PCR using primers NpF4142-5’-F, NpF4142-5’-R, NpF4142-3’-F and NpF4142-3’-R and cloned into pRL278^28^ as a BamHI-SacI fragment using restriction sites introduced on the primers.

To construct plasmid pGAP109 for replacement of the chromosomal allele of *fraI* with a C-terminal *gfpuv*-tagged variant, approximately 900 bp of DNA downstream of the stop codon were amplified via PCR using primers NpF4142-gfp-3’-F and NpF4142-3’-R and cloned into pSCR569 ^29^, as an SpeI-SacI fragment using restriction sites introduced on the primers. The coding region and approximately 900 bp of DNA upstream of the start codon were then amplified via PCR using primers NpF4142-5’-F and NpF4142-gfp-5’-R and cloned into this plasmid as a BamHI-SmaI fragment using restriction sites introduced on the primers.

Generation of transposon mutants and identification of transposon insertion sites was performed as previously described ^30^ using plasmid pRL1063a^31^. Gene deletions and allelic replacements were performed as previously described^32^ with *N. punctiforme* cultures supplemented with 4 mM sucralose to inhibit hormogonium development and enhance conjugation efficiency ^30,33^. To construct UCCS103, plasmid pDDR558 was introduced into wild type *N. punctiforme*. To create UCCS113, plasmid pGAP109 was introduced into UCCS103.

### Motility and phototaxis assays

Plate and time-lapse motility and phototaxis assays were performed as previously described^34^ and imaged using a Leica SD9 dissecting microscope equipped with a Flexcam C3 camera and controlled by Leica X LAS X software. The students T-test was applied to determine statistical significance in colony spreading assays.

### Light- and fluorescence microscopy

A Leica DM2500 B microscope equipped with a Leica DFC420C camera, or an Evos M5000 microscope was employed for light microscopy. Fluorescence microscopy utilized a Leica DM5500 B microscope with a 100x/1.3 oil objective lens, connected to a Leica DFC360FX camera. GFP and chlorophyll fluorescence were induced using BP470/40 nm or BP535/50 nm filters, and emission was detected with BP525/50 nm or BP610/75 nm filters, respectively.

Septal visualization was performed using Vancomycin-FL (Bodipy-FL, Invitrogen) fluorescent stains that specifically stains septal peptidoglycan by the previously described procedure^35^.

### FRAP and CCCP-FRAP

A three-day-old liquid culture of vegetative filaments from both the *N. punctiforme* wild type and the Δ*fraI* mutant strains was utilized for CCCP-FRAP, following previously established protocols ^8,36^. Briefly, 2 ml of culture underwent triple washing with BG11 medium and were then resuspended in 500 μL of medium. Subsequently, the cells were incubated with 10 – 15 μL of 1 mg/ml calcein in DMSO (Sigma-Aldrich) for 90 minutes in the dark at 28 °C with shaking. Following this incubation period, the cells underwent three additional washes with BG11 medium and were then either resolved in medium alone or in medium containing CCCP (from a 50 mM stock solution in DMSO), achieving a final concentration of 50 μM in the culture. After another 90-minute incubation with shaking at 28 °C in the dark, the filaments were spotted onto BG11 agar plates (1%).

The FRAP procedure was conducted by employing a laser with 488 nm excitation at 0.2% intensity on a Zeiss LSM 800 confocal microscope equipped with a 63x/1.4 oil-immersion objective. The ZEN 2.9 (blue edition) software was used for the experiment. Simultaneous detection of calcein fluorescence emission (400–530 nm) and chlorophyll autofluorescence emission (650–700 nm) was carried out. To bleach a specific cell, the laser intensity was briefly increased to 3.5% after capturing 5 pre-bleached images. Subsequently, fluorescence recovery in the bleached cell was observed at 1-second intervals over a period of 30– 120 seconds. Image analysis was performed using ImageJ (version 1.51j) and GraphPad Prism 10, as detailed in previous descriptions.

For hormogonia FRAP, a log-phase culture grown with 4 mM sucralose in BG11 medium was washed twice with BG11 for hormogonia induction. After 18-20 hours following hormogonia induction the cells were harvested (3-5 ml) and used for FRAP as described above. The stained cells (25 μL) were put onto a plain microscopic slide and covered with a cover slip. The measurement was only conducted after the hormogonia ceased movement upon becoming trapped between coverslip and slide due to evaporation.

### Akinete induction

Akinetes were induced essentially as described^37^. In short, liquid cultures of WT and NpF4142 mutant were grown in BG11 medium with nitrate until early log-phase. Akinete formation was induced by washing the culture twice followed by inoculation in phosphate-free BG11 medium and incubation with low shaking (50 rpm) for 4 weeks.

### Isolation of septal peptidoglycan and transmission electron microscopy

Septal peptidoglycan was isolated as described^36^. First, vegetative filaments of *Nostoc punctiforme* WT and Δ*fraI* grew for 5 days in liquid BG11 medium supplemented with 4 mM sucralose to supress hormogonia development. For hormogonia induction, a log-phase culture was induced by washing twice with BG11 medium and incubated in fresh BG11 medium for 20 h. The vegetative and hormogonia cultures were washed and resuspended in 700 μL 0.1 M Tris-HCl pH 6. Briefly, cells were sonicated (Branson Sonier 250), then boiled in SDS for 30 min. Then, the samples were sonified in a water bath for 30 min, followed by washing and incubation in 50 mM Na_3_PO_4_ pH 6.8 with 300 μf of α-chymotrypsin. The chymotrypsin was readded and the next day the cells were again sonicated for 30-60 s. Purified septa (10 μL) were added onto an UV-irradiated (16 h) formvar/carbon film-coated copper grid (Electron Microscopy Sciences) for 30 min, stained with 1% (w/v) uranyl acetate and imaged with a Philips Tecnai10 electron microscope at 80 kV equipped with a Rio Camera (Gatan). Images were analysed using ImageJ 2.9. to calculate the number of nanopores per septa and the septum’s diameter.

### Transmission electron microscopy of ultrathin sections

To examine hormogonia via transmission electron microscopy, we induced their formation by washing of the cells previously cultivated in AA/4 media with 2.5 mM of sucralose. After 20 hours, hormogonia were harvested by centrifugation and ultrathin sections were prepared following standard procedure^38^. Prior to sectioning, the cells underwent fixation with glutaraldehyde and potassium permanganate. Subsequently, the ultrathin sections were stained with uranyl acetate and lead citrate. Finally, the samples were analysed using a Philips Tecnai10 microscope operating at 80 kV.

### Immunolocalization of AmiC2

Immunolocalization of AmiC2 in *Nostoc* strains was performed based on two protocols^17,39^. *Nostoc punctiforme* cultures were cultivated in AA/4 medium with supplemented NaNO_3_, KNO_3_ and 2.5 mM sucralose, washed once and resuspended in 1 mL PBS. The cells were fixed with 4% paraformaldehyde PBS (137 mM NaCl, 2.7 mM KCl, 10 mM Na_2_HPO_4_, 1.8 mM KH_2_PO_4_, pH 7.4) on ice for one hour, followed by three rounds of washing with PBS and final incubation in GTE buffer (50 mM glucose, 25 mM Tris-Cl, pH 8.0, 10 mM EDTA). Then, 200 μL of the cell suspension were put onto Epredia™TM Polysine adhesion slides (Epredia, Netherlands) and let to dry. The cells were then fixed with -20 °C methanol for 5 min followed by incubation with -20 °C acetone for 30 s. Cells were dried at RT and rehydrated with 200 μL PBS for 5 min. After blocking with 200 μL 2 % w/v BSA in PBS for 30 min, cells were incubated overnight in a wet chamber at 4 °C with 1:5 anti-AmiC1 ^17^ in 2 % w/v BSA BSA-PBS. The slides were washed three times with PBS, followed by a two-hour incubation at room temperature in darkness with FITC-coupled α-rabbit antibodies (diluted at 1:200 in BSA-PBS from Sigma Aldrich). After subsequent washing and drying, a drop of Vectashield Mounting Medium H-1200 (Vector Laboratories, USA) was added. The preparation was covered with a coverslip. Fluorescence imaging was conducted with a DM5500B Leica microscope and a DFC360FX monochrome camera. Autofluorescence was measured as mentioned above, while FITC-fluorescence was captured using a BP470/40 nm excitation filter and a BP525/50 nm emission filter. Z-stacks with 0.20 μm intervals were acquired and 3D-deconvolution using Leica ASF built-in function was performed.

### Test on significance

Statistical analysis was performed with GraphPad Prism version 10. Comparison of one group with multiple other groups was performed via Welch ANOVA test followed by Brown-Forsythe followed by Dunnett’s T3 multiple comparison test. Significance *P* is indicated with asterisks: *: *P*≤0.05; **: *P*≤0.01; ***: *P*≤0.001; ****: *P*≤0.0001; ns (not significant): *P*>0.05.

## Results and Discussion

### *fraI* deletion affects hormogonia motility, filament length and diazotrophy

Using a transposon mutagenic screen^30^ we identified a mutant strain with reduced motility containing a transposon insertion in open reading frame Npun_F4142 (Table S1). Its homologue in *Nostoc sp*. 7120 that we previously named FraI^40^ (Kieninger 2023) was shown to be highly abundant in a co-immunoprecipitation experiment against septal junction proteins FraD and SepN ^36^ and displayed a fragmentation phenotype^40^. Based on this, and the data presented below, we have designated the gene name for Npun_F4142 as *fraI*. To confirm that the transposon insertion in *fraI* was responsible for the motility defect, a strain with an in-frame deletion of *fraI* was constructed (Δ*fraI*). Compared to the wild type, the Δ*fraI* strain exhibited a significant reduction in colony spreading in plate motility assays (Fig. 1A). Moreover, there was a qualitative difference in the appearance of motile colonies for the Δ*fraI* strain, with filaments spreading in collective masses rather than dispersed filaments as observed in the wild type (Fig. 1A). In time-lapse motility assays of individual hormogonium filaments (Video S1) the Δ*fraI* strain exhibited motility of individual filaments, but the filaments were much shorter than those of the wild type, and many of the shortest filaments were non-motile.

**Figure 1.**
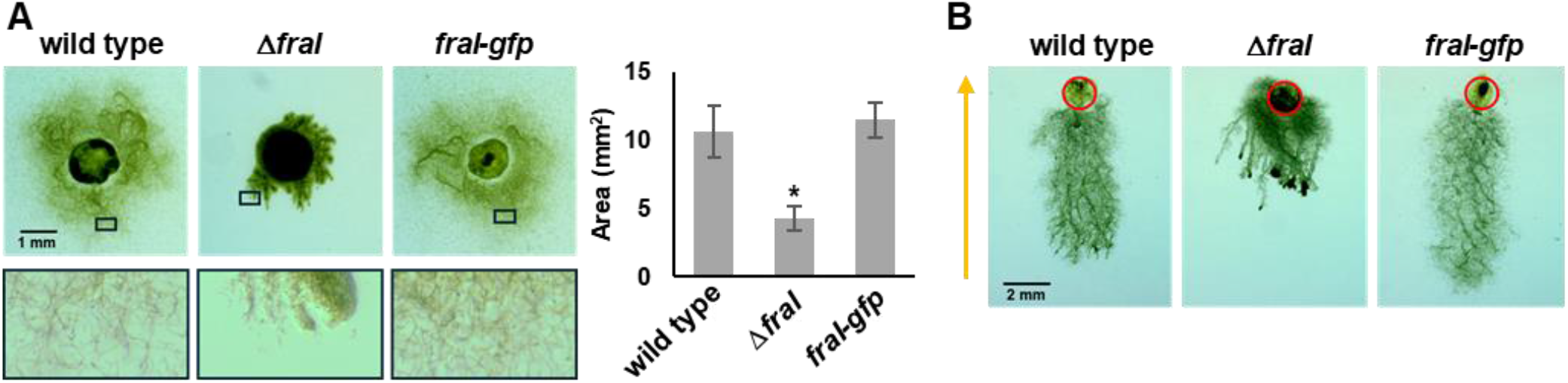
Deletion of *fraI* affects motility. **(A)** Images of plate motility assays (left) and quantification of colony spreading (right) of strains as indicated. For images on left, black boxes in top panel correspond to closeup images of colony periphery shown in bottom panel. For quantification, n=3, error bars =+/- 1 S.D., *=p-value <0.01 vs wild type. **(B)** Images of phototaxis assays for strains as indicated. Red circle indicates the initial site of colony at beginning of assay. Direction of light indicated with arrow.

In an attempt to visualize the localization of FraI, a *fraI-gfp* allele was re-introduced at the native chromosomal locus of the Δ*fraI* strain. Introduction of *fraI-gfp* into the deletion strain restored wild-type motility indicating this allele is functional (Fig. 1A), and further demonstrating that deletion of *fraI* was responsible for the observed phenotype. However, no fluorescence signal could be visualized, indicating that the abundance of FraI-GFP in the cell may be below the limit of detection, or that fusion of GFP to the C-terminus of FraI may affect GFP folding or stability. Attempts were made to introduce either the native, or a *gfp*-tagged allele of *fraI* into the deletion strain on a replicative shuttle vector, but in both cases these efforts failed to yield viable colonies, possibly implying that increased expression of *fraI* from a multicopy plasmid is lethal in *N. punctiforme*. Given the motility defect in the Δ*fraI* strain, its capacity for phototaxis was also investigated (Fig. 1B). Although migration was reduced compared to the wild type, the Δ*fraI* strain exhibited directional movement in response to light, indicating the mutant strain retains the ability to perceive and respond to light signals.

In liquid cultures supplemented with ammonia, the Δ*fraI* strain was more dispersed than the wild type (Fig. 2A). Under these standard growth conditions, the wild-type strain produces a mixture of vegetative and hormogonium filaments (Fig. 2B). The Δ*fraI* strain also produced a mixture of both filament types, but in general, the filaments were much shorter (Fig. 2B). Moreover, we frequently observed what appeared to be hybrid filaments containing a mixture of vegetative and hormogonium cell types in the same filament (Fig. 2B). Although difficult to quantify given the tendency of *N. punctiforme* to aggregate into dense clumps, this phenotype was never observed in the wild-type strain, nor have we ever observed such a phenotype in previous studies with either the wild type or any of the mutants we have characterized. These results indicate that while *fraI* is dispensable for hormogonium development and motility, it may be critical for cells within a filament to synchronize hormogonium differentiation.

**Figure 2.**
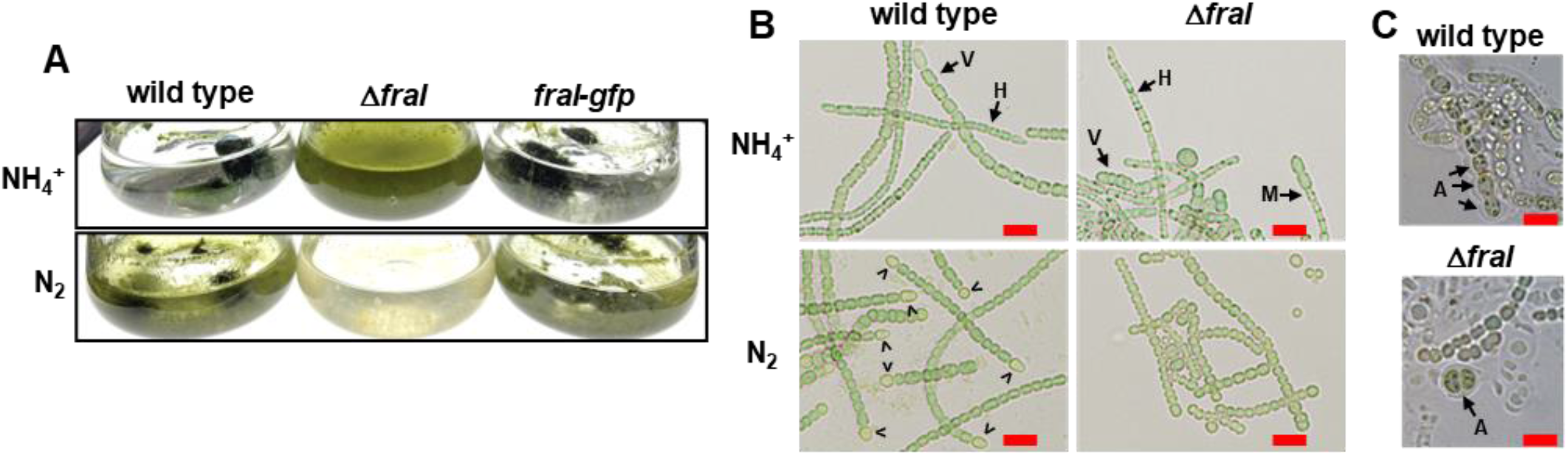
Influence of *fraI* on diazotrophy and development. **(A)** Images of cultures (strains as indicated) in presence (NH_4+_) or absence (N_2_) of a source of fixed nitrogen. **(B)** Light micrographs of individual filaments (strains as indicated) in presence (NH_4+_) or absence (N_2_) of a source of fixed nitrogen. V = vegetative filament, H = hormogonia, M = mixed filament containing both hormogonium and vegetative cells, > = heterocysts. Bar = 10 um. **(C)** Light micrographs (strains as indicated) of filaments following induction of akinetes by removal of phosphate from the growth medium. A = akinete, Bar = 10 um.

In liquid cultures lacking a source of fixed nitrogen, conditions where the wild type develops nitrogen-fixing heterocysts to support growth on N_2_, the Δ*fraI* strain became chlorotic and failed to grow (Fig. 2A). Under these conditions the fragmentation phenotype became more severe, and there was no obvious indication that morphologically distinct heterocysts were present (Fig. 2B). As with motility, the introduction of *fraI-gfp* into the deletion strain restored diazotrophy (Fig. 2A). Upon phosphate starvation the Δ*fraI* strain was able to develop akinetes although of aberrant shape and not in consecutive manner when compared to the WT^37^ (Fig. 2C). Collectively, the absence of heterocysts and fragmentation phenotype of the Δ*fraI* mutant resembles that reported for mutants in other genes known to be involved in formation of the septal junctions, most of which have been characterized primarily in the closely related filamentous cyanobacterium *Nostoc* sp. PCC 7120.

The reduced motility phenotype motivated us to examine the ultrastructure of hormogonia filaments using transmission electron microscopy (TEM). In Δ*fraI* we could observe short hormogonia that were completely lacking the surrounding exopolysaccharide sheath (Fig. 3), which may be the cause of the of observed reduced motility (Fig. 1). In addition, no septal cell-cell connections were visible in Δ*fraI* hormogonia (Fig. 3). Furthermore, in contrast to the WT, the thylakoid membranes were aberrantly arranged. This could be explained by the presence of numerous glycogen granula, which may disturb the organization of the membranes.

**Figure 3.**
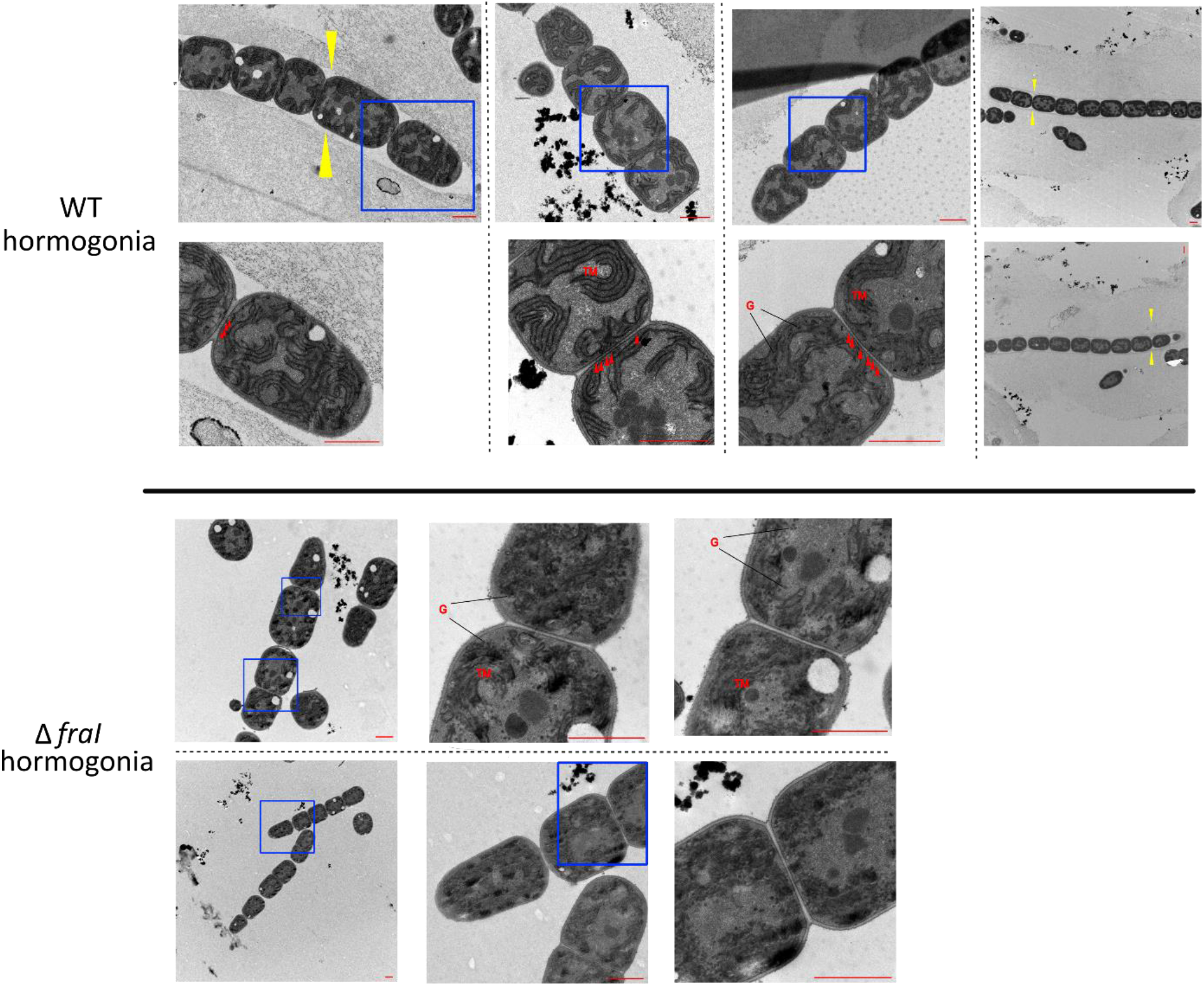
The hormogonia of Δ*fraI* have no cell-cell connections and are lacking the HPS. After induction of hormogonia, ultra-thin sections were prepared and the ultrastructure investigated by transmission electron microscopy. Shown are representative electron micrographs of each strain. Blue rectangles signify the zoomed region shown below (WT) or at the right (Δ*fraI*). Red arrows point to the cell-cell junctions visible in the WT. Yellow arrows point to the exopolysaccharide sheath present only in the WT hormogonia. G – glycogen granules, TM – thylakoid membrane. Scale bar – 1 μm.

### FraI is related to known septal proteins

To further explore the relationship between FraI and septal junctions, we investigated the expression pattern of *fraI* as well as the orthologs of other septal junction genes during hormogonium and heterocyst development in *N. punctiforme* using previously published transcriptomic data sets^29,41,42^. Additionally, the evolutionary co-occurrence of this gene set in cyanobacteria was determined. During hormogonium development all the identified septal junction genes are transcriptionally upregulated in wild-type *N. punctiforme*, with expression typically peaking between 6 to 12 h post induction (Fig. S1A). This observation is consistent with the fact that developing hormogonia undergo a round of cell division, necessitating the construction of large numbers of new septal junctions at the new septa. For most of these genes, enhanced transcription in hormogonia is independent of the hormogonium sigma factor cascade^41^, as deletion of *sigJ, sigC*, or *sigF* has no obvious effect on expression. In contrast, upregulation of *fraI, sepI* and *sjcF1* in hormogonia is dependent on both *sigJ* and *sigC*. Given that expression of *sigC* is also dependent on *sigJ*, these results imply that these three genes are most directly regulated by *sigC*. Transcriptional start site mapping via Cappable seq (CAPseq) indicated the presence of two transcriptional start sites (-265 bp and -200 bp of the *fraI* start codon) in the *fraI* promoter region, with the more distal closely corresponding to total read coverage from RNAseq data (Fig. S1B). However, this TSS only showed a very minor decrease in abundance in the Δ*sigC* strain. In developing heterocysts there was less pronounced upregulation of this gene set, with only a few genes displaying marked upregulation (Fig. S1A). Co-occurrence analysis indicated that FraI was found almost exclusively in heterocyst-forming cyanobacteria, similar to the conservation pattern of SepN, FraD, and FraC, although orthologs of these genes are found occasionally in non-heterocyst forming filamentous cyanobacteria (Fig. S2). Overall, the expression and co-occurrence patterns of *fraI* in comparison to other septal junction genes support a role for *fraI* in the formation of septal junctions.

### The Δ*fraI* mutant is severely impaired in cell-cell communication

The relation to septal junction proteins and the Δ*fraI* phenotype ^10,11,43^, prompted us to investigate cell-cell communication and septal junction gating ability in the Δ*fraI* mutant of *N. punctiforme*^8,36^. The study of cell-cell communication was performed by FRAP experiments

^10,44^, where filaments were first stained with a fluorescent dye, subsequently, a single cell is laser-bleached, and finally the recovery of fluorescence, resulting from the diffusion of the dye through septal junctions, was monitored over time^10,44^. Compared to the wild type of *N. punctiforme*, the vegetative cells of the Δ*fraI* mutant were highly impaired in cell-cell communication, with 39% of the cells showing no communication at all, and 57% of the cells showing a very low communication rate (R < 0.02 s^-1^) (Fig. 4A). To investigate gating ability of septal junctions, the cells were treated with the protonophore CCCP which induces closure of SJ^8,36^. Incubation with CCCP caused almost complete halt of communication (86% of the cells) in the WT and in the Δ*fraI* mutant (77%) (Fig. 4A). The recovery rates were calculated for “normal recovery” and “slow increase” cells. Here, recovery rates in vegetative filaments are 0.053 s^-1^ and 0.012 s^-1^ for WT and Δ*fraI* mutant respectively.

**Figure 4:**
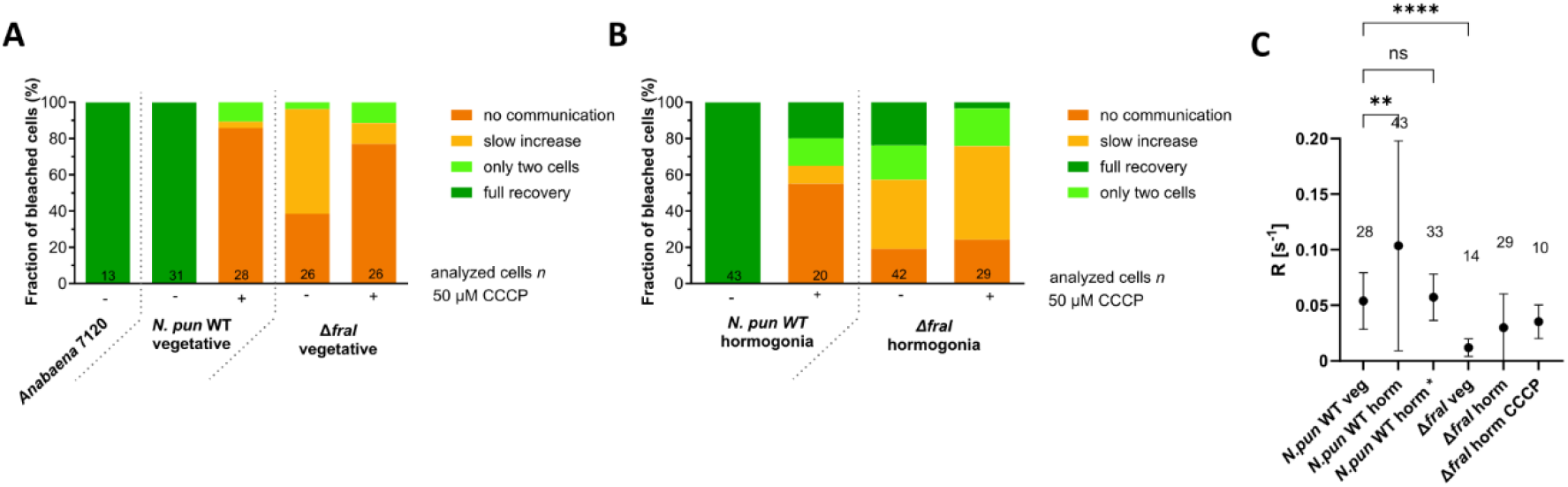
The Δ*fraI* mutant is highly impaired in cell-cell communication in both vegetative filaments and hormogonia. CCCP-FRAP analysis of *Anabaena* PCC 7120, *N. punctiforme* WT (*N. pun* WT) and Δ*fraI* mutant vegetative filaments **(A)** and hormogonia **(B)**. FRAP-responses of calcein-stained untreated or CCCP-treated (90 min, 50 μm) were assigned to one of four groups indicated by the color scheme. The numbers within the bars indicate the number of analyzed cells (n) from different filaments. Recovery rates (*R*) of communicating cells **(C)**. Mean is shown. One-way ANOVA followed by Brown-Forsythe multiple comparison test with the WT was performed. The numbers above the mean indicate the number of cells used for calculation of *R. horm –* hormogonia. *N*.*pun WT horm \** excludes cells in hormogonia of the WT with measured R > 0.19 (see main text for explanation). Cumulated results from at least two independent cultures are shown. **, p<0.01****, p<0.0001; ns, not significant

Since FRAP experiments haven’t yet been described in hormogonia, we wanted to measure the rate of communication between the cells in hormogonia and the cell’s ability to close septal junctions upon treatment with the protonophore CCCP (Fig. 4B,C). On average, WT hormogonia showed much higher rates of communication (R = 0.104 s^-1^) compared to vegetative cells (R = 0.054 s^-1^) (Fig. 4C), with a fraction of the cells (23%) showing very high recovery rates (R > 0.19 s^-1^). Many cells undergo cell-division more or less simultaneously during hormogonia development and hence in many cases the new septum may have not yet completely closed, explaining the large number of cells with such unusual high communication rates in FRAP analysis ^45,46^. This idea is also supported by the fact that only 55% of the WT hormogonia cells completely stopped communication after CCCP treatment, in contrast to 86% among vegetative cells (Fig. 4B). Also, 76% of Δ*fraI* mutant hormogonia cells showed a communicating phenotype, compared to 61% of the vegetative cells, because, like in the WT, cell division was not yet completed and the fluorescent dye could easily diffuse between the future daughter cells (Fig. 4B).

The inability of the Δ*fraI* mutant to differentiate heterocysts can be explained by the lack of cell-cell communication. In contrast, the Δ*fraI* mutant can still differentiate akinetes despite decreased cell-cell communication”. This is in line with our previous observation with the related species *Trichormus* (*Anabaena*) *variabilis* ATCC29413. There, we showed that during akinete formation molecular transfer decreases and seems not to play a role in this spore differentiation process^47^.

#### The Δ*fraI* mutant lacks the septal nanopore array

To investigate whether aberrant nanopore array formation is causing the impaired communication in the Δ*fraI* mutant, the septal peptidoglycan of vegetative cells and hormogonia of the WT and the mutant were isolated and analyzed by TEM. The Δ*fraI* mutant showed very few or almost no nanopores, both in PG from vegetative filaments as well as from hormogonia (Fig. 5A,B). Interestingly, the Δ*fraI* mutant also had a significantly larger septa diameter of vegetative filaments compared to the WT (Fig. 5B). A similar phenotype, namely the lack of nanopores in septa and enlarged septa was previously observed for an *amiC2* mutant of *N. punctiforme*^35^. Interestingly, the *amiC2* mutant displayed delocalized septa leading to a severe morphological phenotype^12^, which is not observed for the Δ*fraI* mutant using Vancomycin-FL (Figure S4). In contrast, the pores in the lateral wall just around the septal discs, were present in the Δ*fraI* mutant and the WT (Fig. 5A,B), implying that FraI is not involved in formation of these lateral pores. In the center of the septal discs of the WT and Δ*fraI* mutant hormogonia, we could occasionally observe few very large nanopores (Figure S5), which may also explain higher rates of communication measured in some hormogonia cells (see above).

**Figure 5:**
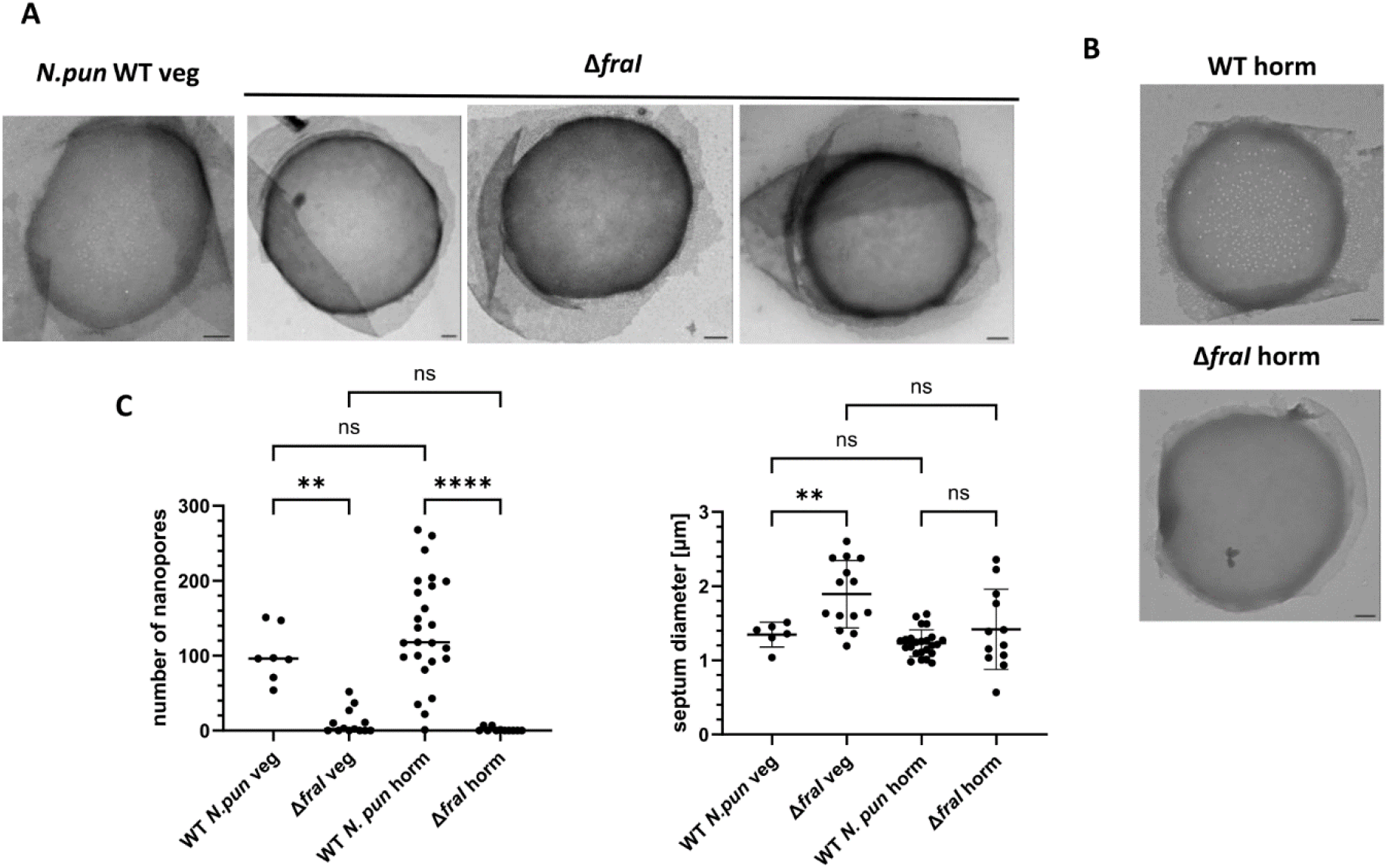
The Δ*fraI* mutant completely lacks the septal nanopore array and has greater septa diameter. Septal peptidoglycan in *N. punctiforme* WT and Δ*fraI* mutant vegetative filaments **(A)** and hormogonia **(B)**. Representative TEM images are shown. Scale bar – 200 nm. **(C)** Number of nanopores and septum diameter of *N. punctiforme* vegetative and hormogonia cells in WT and Δ*fraI* mutant, which has very few or no nanopores and a greater septa diameter. One-way ANOVA followed by Brown-Forsythe multiple comparison test. Median is shown. p<0.05 ** p<0.01, **** p<0.0001, ns p>0.05.

Despite almost complete lack of nanopores, the AmiC2 protein could be detected in the septa of Δ*fraI* mutant using anti-AmiC2 antibodies (Figure S3). Hence, FraI is unlikely to be involved in septal localization of AmiC2. Moreover, FraI is predicted to have three transmembrane helixes with a cytoplasmic C-terminus and an N-terminal signal sequence of just 20 amino acids residing in the periplasm (Phobius), which makes a direct protein-protein interaction with the periplasmic amidase unlikely. However, at the moment we cannot exclude that a regulatory protein, like the LytM factor, is influenced by FraI. In line with the idea of a FraI-AmiC2 relation is the fact that *fraI* expression is the highest during hormogonia development (Figure S1), when many cells are undergoing cell division and the need for AmiC2 activity is the highest^48^. The lethality of overexpressed FraI (see above), is another indication that this protein is involved in regulation of the cell wall lytic enzyme AmiC, since the over-expression of the *amiC* orthologues in *Nostoc* sp. PCC 7120 was also lethal^13^.

It is interesting that the impaired cell-cell communication rate in the Δ*fraI* mutant did not prevent hormogonia development. However, the motility of the hormogonia was reduced (Fig. 1A). Nevertheless, the Δ*fraI* mutant can move towards the source of light, indicating that cell-cell communication through the septal junctions is not crucial for hormogonia motility and synchronized movement along the filament in response to a positive external signal. Additionally, the Δ*fraI* mutant was able to change the direction of movement following a change of light conditions (data not shown). Therefore, there may exist another way of communication, perhaps through the common periplasm, which enables synchronized movement of the filament. As an alternative explanation, each single cell may decide the direction of the movement based on the sensing of external factors such as light and chemoattractant gradient. It has already been speculated that a change in an electromotive force may be a signal for translocation of HmpF protein of the pili machinery to the opposite side of the cell, which may be sensed by each cell in hormogonia^49^. Furthermore, the observed reduction in motility in the Δ*fraI* mutant could be attributed to the diminished presence of the exopolysaccharide sheath surrounding hormogonia filaments (Fig. 3). Previous studies have demonstrated reductions in motility in mutants with impaired exopolysaccharide synthesis and export pathways ^32,34^. Why the absence of *fraI* gene leads to lower presence of hormogonia-specific exopolysaccharide remains unknown. It has been speculated that HPS is exported through the periplasm close to the cell junctions in hormogonia^50^.

Alternatively, despite being significantly impaired in communication, 81% of Δ*fraI* hormogonia cells (Fig. 4) still exhibit some communication, possibly facilitated by the presence of remaining nanopores or incomplete septa formation. This residual communication may still be sufficient for the transport of signal necessary for the uniform polar distribution of pili machinery enabling directionality of movement along the filament in Δ*fraI* mutant.

## Conclusion

In this study we discovered FraI, a new player in cell-cell communication and differentiation. Mutation of *fraI* leads to a low-communicating phenotype in both vegetative cells and hormogonia of *Nostoc punctiforme*, which can be explained by a nearly complete absence of nanopores in the septa of neighboring cells. Despite the low levels of communication, the Δ*fraI* strain forms motile hormogonia capable of synchronized movement, albeit in a different way than the WT. The filaments also preform phototaxis, suggesting that motility and directionality of movement do not require communication through the septal junctions. Lower motility observed in Δ*fraI* is more likely to be explained by the absence of the HPS sheath. In contrast, the presence of filaments containing a combination of vegetative and hormogonium cells in the Δ*fraI* mutant, a phenotype never observed in the wild type, implies that communication through the septal junctions may play a critical role in coordinating the development of hormogonia. As in other mutants of septal nanopore array formation, severely impaired cell-cell communication of the Δ*fraI* mutant is affecting heterocyst differentiation and function and hence diazotrophic growth. Nonetheless, akinete development is still possible in the Δ*fraI* mutant. FraI may play a role in indirectly controlling amidase activity through an unknown mechanism. Additional experiments focusing on the FraI protein will be necessary to support this claim. Finally, characterization of orthologues of known septal junction proteins from *Nostoc* sp. 7120 in *N. punctiforme* will provide additional understanding of the role of septal junction-mediated communication in the process of hormogonium development and motility.

## Author contribution

AJ: designed and performed experiments, analyzed and interpreted data, drafted the work and wrote the manuscript.

CM: performed TEM experiments and sample preparation for ultra-thin sections.

DDR: designed and performed experiments, analyzed and interpreted data, made considerable contributions to the design of the study, interpreted data, wrote the manuscript.

GAP: designed and performed experiments, analyzed and interpreted data

IM: designed and supervised the research in Tübingen, made considerable contributions to the design of the study, interpreted data, revised the manuscript critically.

All authors approved the final manuscript.

## Data Availability

Data available from authors upon reasonable request.

