## Supplemental Material for "The role of FraI in cell-cell communication and differentiation in the hormogonia-forming cyanobacterium Nostoc punctiforme"

**Table S1**. - Strains, plasmids and primers used in this study.

| **strain name** | **description** |
|---|---|
| *N. punctiforme* ATCC29133 | wild type |
| UCCS 103 | In-frame deletion of *fraI* (Δ*fraI*) |
| UCCS 113 | Native *fraI* allele replaced with *fraI-gfpuv* |
| UCTN1 | Tn5 insertion at 225 bp of fraI coding-region |

| **plasmid name** | **description** |
|---|---|
| pDDR558 | Suicide vector for in-frame deletion of *fraI* (Npun_F4142) |
| pGAP109 | Suicide vector for replacement of native *fraI* allele with *fraI-gfpuv* |

| **Primer name** | **sequence** |
|---|---|
| NpF4142-5'-F | atataggatccGGAAACGCACTGATTGAAAAC |
| NpF4142-5'-R | cactttattacctttgCAATAGAAACATAGATTAACTCTC |
| NpF4142-3'-F | ctatgtttctattgCAAAGGTAATAAAGTGAGAGAG |
| NpF4142-3'-R | atatagagctcAAAGGGTAAACTGCCAGAG |
| NpF4142-gfp-5'-R | atatacccgggACCTTTGAACACGCAAGAAC |
| NpF4142-gfp-3'-F | atataactagtTAAAGTGAGAGAGAAGTCAG |

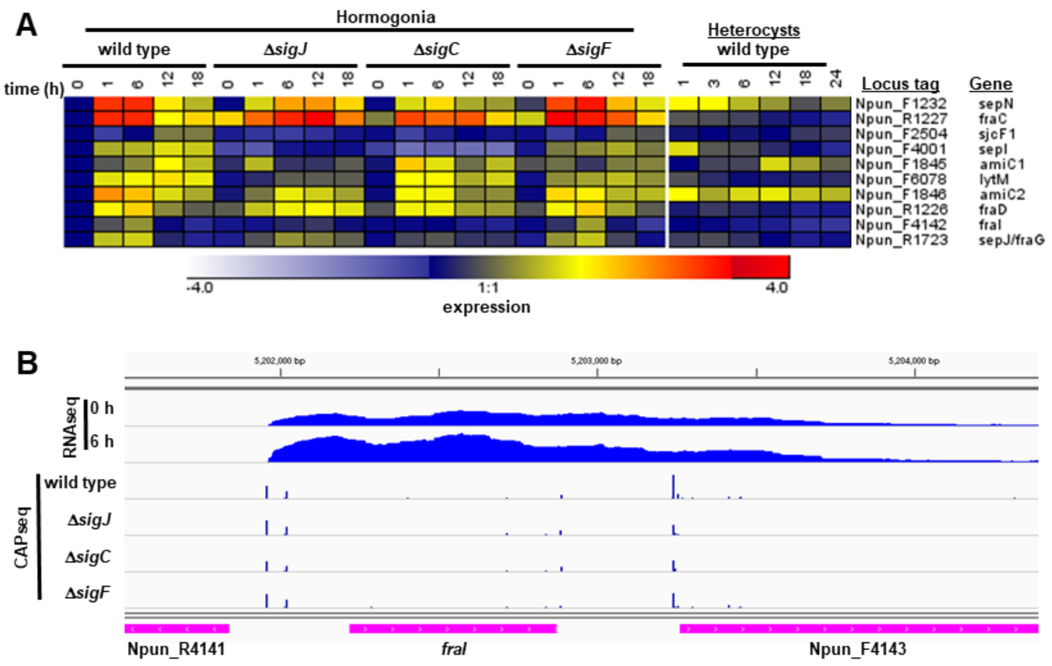


**Figure S1.** Transcription of septal junction genes in developing hormogonia and heterocysts in *N. punctiforme* **(A)** Heat maps depicting the expression of *fraI* and other septal junction genes in developing hormogonia of the wild-type and hormogonium-specific sigma factor mutants 0-18 h post hormogonium induction, or in developing heterocysts 0-120 h post heterocyst induction. Expression = Log2 (experimental strain and time point/wild type t=0). **(B)** Read map coverage of the *fraI* locus from RNAseq (0 or 6 h post hormogonium induction) and Cappable-Seq (6 h post hormogonium induction) data for various strains as indicated.

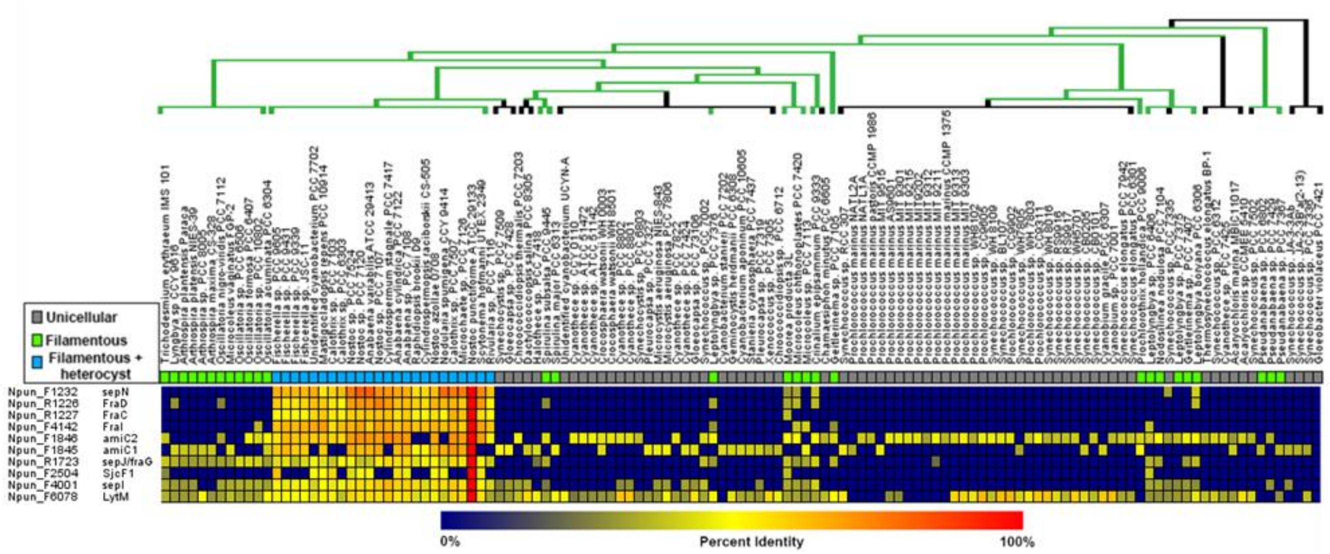


**Figure S2.** Evolutionary conservation of genes encoding septal junction proteins in cyanobacteria. Heat map depicting the percent identity for orthologs of *N. punctiforme* septal junction genes in cyanobacteria, derived from data reported by Cho *et al*.[23]. Species organization and phylogenetic tree based on the phylogeny reported by Shih *et al*.[51] , but depicting the finding, as reported by Schirrmeister *et al*.[52], that most extant cyanobacteria are derived from a filamentous ancestor. For the phylogenetic tree, green = filamentous, black = unicellular.

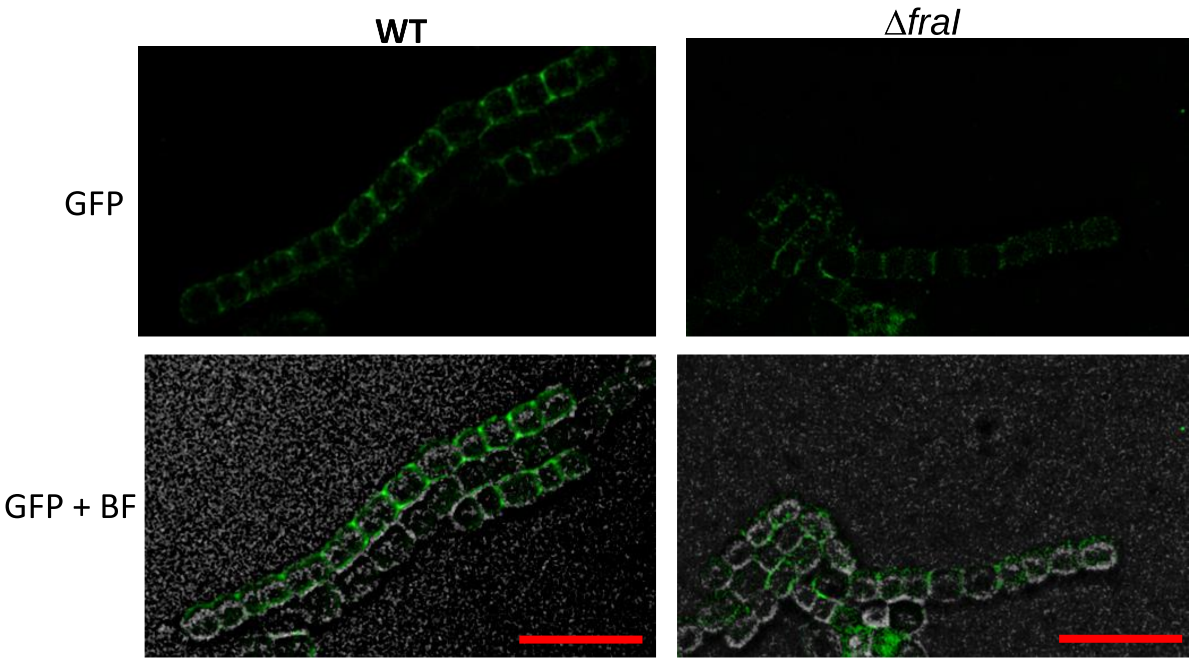


**Figure S3** – Immunolocalization of AmiC2 in *Nostoc punctiforme* WT and Δ*fraI* mutant. Scale bar – 10 µm

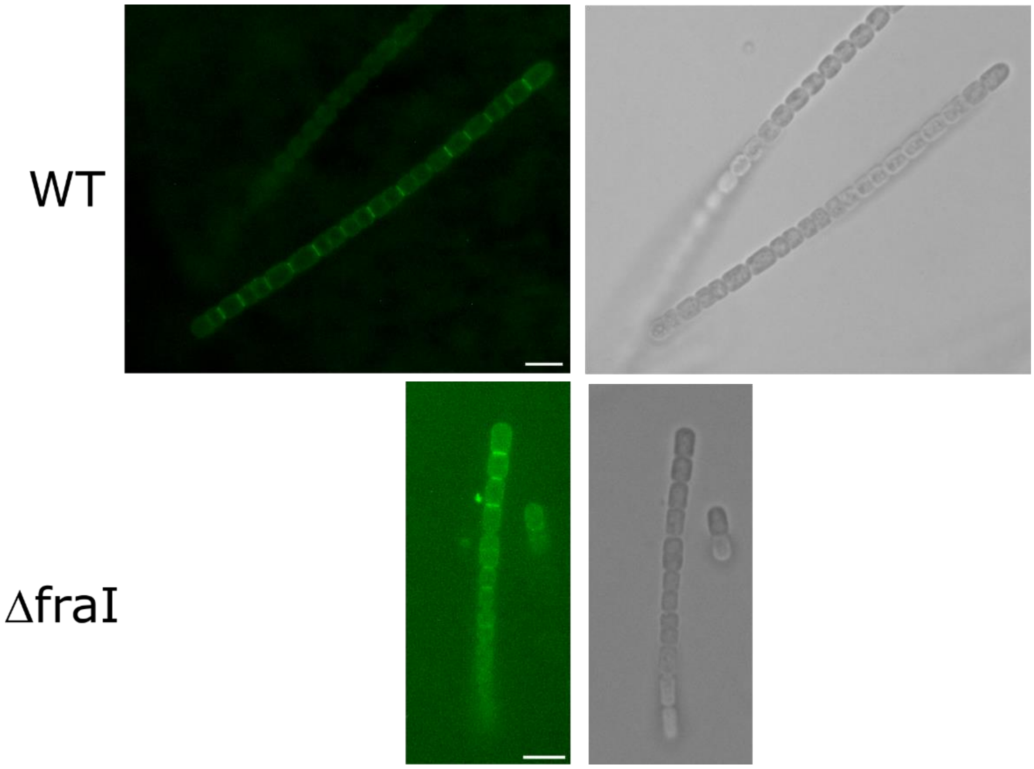


**Figure S4** – Vancomycin-FL staining of hormogonia septal peptidoglycan of WT and Δ*fraI*. Scale bar – 5 µm

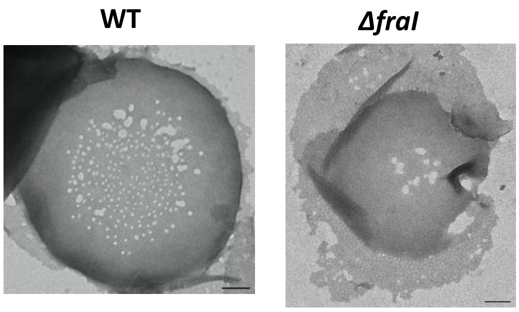


**Figure S5** – Hormogonia septa of WT and Δ*fraI* with large nanopores. Scale bar – 200 nm
